# High resolution protein-protein interaction mapping using all-versus-all sequencing (AVA-seq)

**DOI:** 10.1101/462309

**Authors:** Simeon S. Andrews, Stephanie Schaefer-Ramadan, Nayra M. Al-Thani, Ikhlak Ahmed, Yasmin A. Mohamoud, Joel A. Malek

**Author notes:** These authors contributed equally (Simeon S. Andrews, Stephanie Schaefer-Ramadan). Corresponding author: Correspondence and material requests should be addressed to: Dr. Joel Malek, Weill Cornell Medicine-Qatar, PO Box 24144, Doha, Qatar.

## Abstract

Two-hybrid systems test for protein-protein interactions and can provide important information for genes with unknown function. Despite their success, two-hybrid systems have remained mostly untouched by improvements from next-generation DNA sequencing. Here we present a method for all-versus-all protein interaction mapping (AVA-seq) that utilizes next-generation sequencing to remove multiple bottlenecks of the two-hybrid process. The method allows for high resolution protein-protein interaction mapping of a small set of proteins, or the potential for lower-resolution mapping of entire proteomes. Features of the system include open-reading frame selection to improve efficiency, high bacterial transformation efficiency, a convergent fusion vector to allow paired-end sequencing of interactors, and the use of protein fragments rather than full-length genes to better resolve specific protein contact points. We demonstrate the system’s strengths and limitations on a set of proteins known to interact in humans and provide a framework for future large-scale projects.

## Introduction

Methods of studying protein-protein interactions (PPIs) can broadly be categorized as one-vs-one or one-vs-all studies. The goal of this study is to develop and apply a novel methodology that allows screening in an all-vs-all fashion to compare complex libraries. Here, we have applied our method to define the interacting regions of a set of human proteins with high resolution.

Yeast- and bacterial-2-hybrid screens and their derivatives have long been a staple of large-scale protein interaction mapping ^1,2^. Though there are multiple forms of two-hybrid systems, the most common approaches fuse a protein of interest (“bait”) to a DNA-binding domain (DBD) and test it against a library of proteins (“prey”) that are fused to a transcription activation domain (AD). If the bait and prey proteins interact, they drive the overproduction of a survival gene by bringing the AD and DBD into proximity (Sup. Fig. 1). The interacting proteins are then identified by sequencing bait and prey DNA from proliferating colonies. In the case of the bacterial-2-hybrid (B2H) system used here, the bait protein is encoded as a C-terminal fusion of the λ-cI protein (DBD), while the prey is encoded as a C-terminal fusion of the RNA polymerase α subunit (AD). If the bait and prey interact, the DBD and AD can jointly drive production of HIS-3, a gene which is essential for growth in histidine-free medium, and which can be inhibited by the small molecule 3-amino-1, 2, 4-triazole (3-AT).

Although 2-hybrid screens have been used for many years, only recently has next-generation sequencing (NGS) been applied to facilitate it ^3–5^. However, all these methods still require the use of one-vs-all methodologies: each bait must be sequentially screened against the entire library. Based on the B2H system by Dove and Hochschild ^2^, we have developed a new method which joins both bait and prey as a convergent-fusion into one plasmid called pAVA. This all-versus-all sequencing (AVA-seq) system is amenable to the power of NGS technology and allows for high resolution mapping of protein interaction sites.

## Results

### AVA-seq Overview

The pAVA system is unique in that a two-hybrid screening of a small subset of proteins or even an entire genome can be completed in a relatively short amount of time and can take advantage of the power of NGS. Figure 1 illustrates the AVA-seq system. First, the genomic DNA is sheared into random fragments and is size selected (Fig. 1a). These fragments are then ligated into two different pBORF vectors (pBORF-DBD and pBORF-AD) to allow for open-reading frame (ORF) filtering using carbenicillin (a more stable alternatively to ampicillin) (Fig. 1b). The ORF filtered fragments are amplified with primers allowing tail-to-tail orientation with a linker region with stop codons in all six frames (Fig. 1c) and fused using overlap extension PCR (Fig. 1e). Amplification of the fused fragments allow for cloning into the convergent-fusion vector pAVA that contains the DBD and AD central to the two-hybrid system (Fig. 1f). This is followed by liquid growth in non-selective or selective conditions (Fig. 1g) where interacting fragments have a growth advantage and become a larger fraction of the population. These fragments undergo paired-end sequencing in batches of several million to identify changes in the population of paired-fragment that signify interactors (Fig. 1h). This allows for high resolution snapshots of the interacting domains.

**Figure 1.**
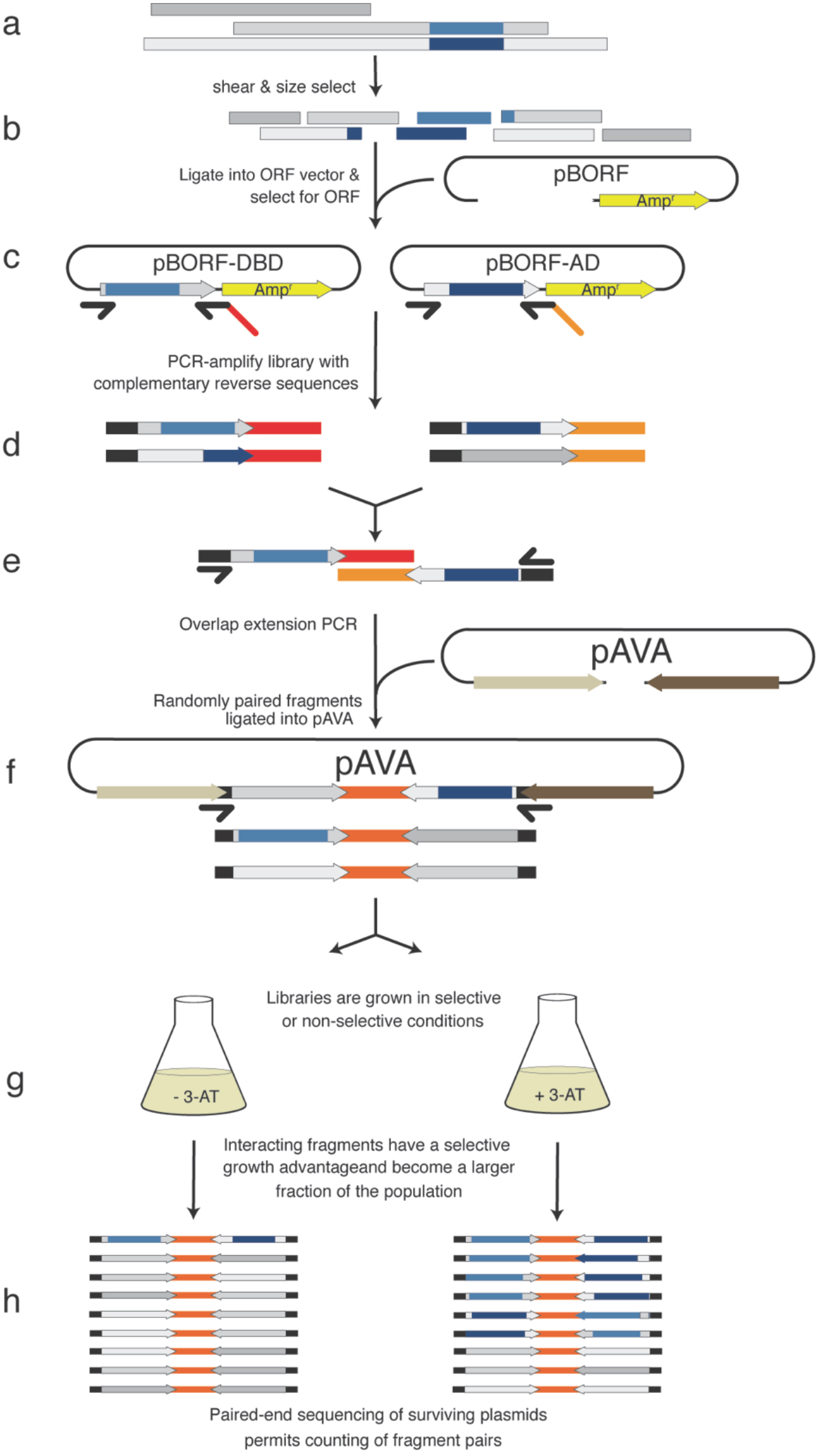
The AVA-seq workflow. To screen a library of proteins (interacting regions colored darker), (**a**) the proteins are sheared, size selected, and (**b**) cloned upstream of a β-lactamase resistance gene in the pBORF vectors. Growth on carbenicillin-containing medium selects for ORF containing fragments. These can be (**d**) amplified with complementary reverse primers, which allows for (**e**) overlap extension PCR to create a fragment pair for (**f**) cloning into pAVA. (**g**) Liquid culture growth in the presence of 3-AT creates a growth advantage for bacteria carrying interacting pairs. (**h**) Paired-end sequencing of millions of inserts allows counting to determine which pairs have increased as a fraction of the population.

### Convergent-Fusion Method Validation

Initial validation of the pAVA convergent-fusion system was conducted with two positive and three negative controls. Growth (OD_600_) was measured following a 9-hour incubation in minimal media in the presence of 0, 2, or 5 mM 3-AT, a competitive inhibitor of the HIS3 gene ^2^. The pAVA vector with no insert (Fig. 2a and f) has strong growth in the unselected (absence of the DMSO vehicle and 3-AT) and 0 mM 3-AT conditions and minimal growth in the presence of 2 and 5 mM 3-AT. To create a positive control of known interactors, Gal11p and LGF2 yeast protein fragments were amplified from the original B2H ^2^ and inserted into the pAVA vector in convergent orientation. The schematics of the convergent-fusion positive control in both orientations are shown in Fig. 2b and c. Both constructs show robust growth in 2 and 5 mM 3-AT (Fig. 2g and h) irrespective of fragment orientation (adjacent to DBD or AD) which indicate a strong interaction between the Gal11p and LGF2, as expected.

**Figure 2.**
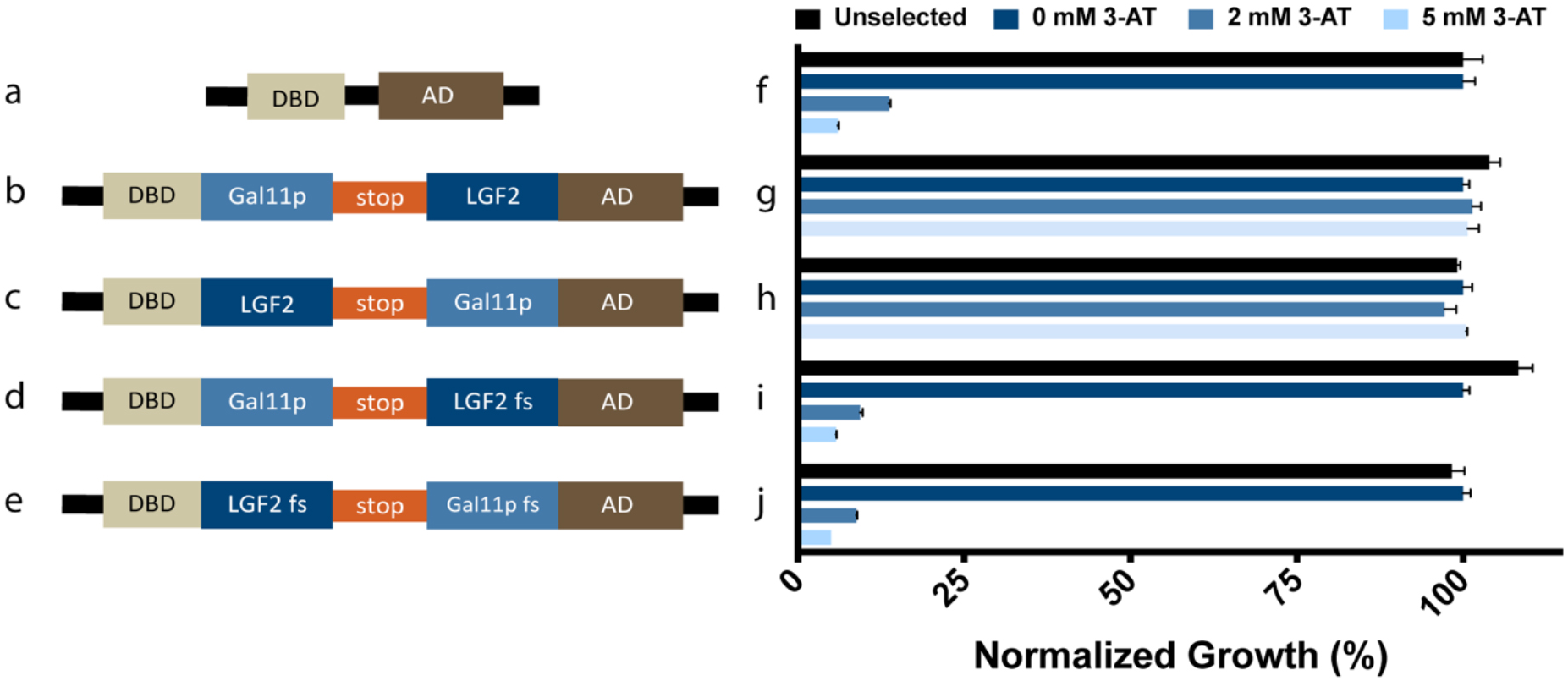
Schematic and growth of controls used in this study. (**a**) pAVA with no DNA insert (negative control), (**b**) pAVA with Gal11p and LGF2 (positive control), (**c**) pAVA with LGF2 and Gal11p (positive control), (**d**) pAVA with Gal11p and LGF2 that has been frame shifted by the insertion of one nucleotide (negative control), (**e**) pAVA with one nucleotide insertion for both LGF2 and Gal11p that results in frameshifts (negative control), (**f**-**j**) competitive inhibition of HIS3 gene using 3-AT. Unselected samples are grown in the absence of 3-AT and DMSO. 0, 2 and 5 mM 3-AT samples have increasing amounts of 3-AT and decreasing amounts of DMSO. Growth charts (**f**-**j**) correspond to schematics (**a**-**e**), respectively. Data represents an average of 3 replicates with error bars representing SEM. The optical density (OD_600_) was normalized to 0 mM growth after 9 hours expression.

To generate negative controls for system validation, nucleotides were inserted at the beginning of the LGF2 sequence (Fig. 2d) or at the beginning of both Gal11p and LFG2 sequences (Fig. 2e) which result in frame-shifts (fs). Both negative controls show diminished growth as the concentration of 3-AT increased, indicating the frame-shifted proteins are not able to overcome the HIS-3 inhibition (Fig. 2i, j). These results indicate that the pAVA system can detect the same protein-protein interaction in both fusion orientations.

### Application to 6 Known Interacting Proteins

After successful validation using yeast Gal11p and LGF2 protein fragments, the pAVA system was tested for high-resolution interaction mapping with 3 pairs of human proteins known to interact *in vivo*. The first pair was transcription factors of the AP-1 complex c-Jun (JUN) and c-Fos (FOS) ^6,7^. The second pair of interacting proteins was cAMP-dependent protein kinase regulatory subunit 2 (PKAR; referred to as PKA in the text) and A-kinase anchoring protein 5 (AKAP5) ^8,9^. The final pair was the human tumor suppressor protein p53 (TP53, referred to as p53 in the text) and the negative regulator of p53 known as murine double minute 2 (MDM2) ^10,11^.

### Open Reading Frame Selection

These 6 human genes were amplified via PCR, pooled together in equimolar amounts and sheared into ~450 base pair fragments (Fig. 1a). A portion of the ligation was subjected to ORF filtering while the remaining portion was not (non-ORF). For ORF selection, the fragments were ligated into both pBORF vectors (AD and DBD) that are modified pBluescript II SK(+) vectors conferring kanamycin (KAN) resistance and containing a split β-lactamase gene. The fragments of interest are cloned between the localization signal and enzyme-coding portion of β-lactamase. Fragments that don’t encode stop codons will permit expression of the β-lactamase gene. Most fragments of 450bp with no stop codon are physiological open-reading frames. pBORF-AD and -DBD differ only in the N-terminal sequence that permits use of non-identical primers in the later step of tail-to-tail fusion PCR. The libraries of fragments in pBORF were challenged with KAN and carbenicillin (CB; β - lactamase expression) using a method similar to Di Niro et al. ^12^. Fragments cloned in the pBORF-DBD and pBORF-AD resulted in 1.3% and 4.8% survival on KAN/CB, respectively, relative to growth on KAN alone. The over 95% drop in clones between the two antibiotics is indication of ORF selection.

Sequencing and analysis of the ORF selected fragments revealed on average 48% of plasmids from pBORF-AD contained in-frame fragments of one of the 6 known proteins used in this study. This compared to approximately 16% in the non-ORF selected libraries in the same vector. ORF selection was less efficient for fragments in pBORF-DBD with approximately 24% being in-frame versus 13% for the non-ORF selected libraries.

### Convergent Fragment Generation and Interaction Screening

The ORF selected fragments in the pBORF-AD and pBORF-DBD were amplified, “stitched” together using overlap extension PCR and inserted into pAVA (Fig. 1d-e). When convergent-fusion fragments were considered, approximately 19% of paired products analyzed contained both DBD and AD fused fragments expressed in-frame with one of the 6 human proteins compared to only 1.5% in the non-ORF selected libraries. ORF selection provides at least a 12-fold improvement in screening efficiency over non-ORF selected libraries, especially given that both fusion fragments are required to be in-frame for interaction. The convergent-fusion fragments in the pAVA plasmid were transformed into the BacterioMatch II Electrocompetent Reporter Cells ^2^ and challenged in triplicate with 0, 2 or 5 mM 3-AT (Fig. 1g) in histidine-free media. The same was process was repeated for the non-ORF fragments.

### Library Construction and Sequencing

Libraries were constructed using the NEBNext Ultra II DNA Library Prep Kit for Illumina according to the manufacturer protocol. A total of 9 libraries, 3 replicates for each 0, 2, and 5 mM 3-AT selection were combined for sequencing using a MiSeq. A total of 6.1 million paired-end sequences were generated for which both the DBD and AD sequences were of high enough quality to allow translation of the fused fragments. This resulted in approximately 680,000 paired reads for each replicate. As control for the selection and sequencing analysis process, the pre-constructed positive control with Gal11p and LFG2 domains (Fig. 2b) was spiked in at low concentration prior to the 3-AT selection.

### Sequence Analysis

Paired-end sequencing reads were translated in-frame with the DBD or AD fragments they were fused to. Sequencing primers sit upstream of the of DBD or AD specific sequence allowing enough sequence (~150bp) downstream to identify the fused fragment and whether the fusion is to DBD or AD. The translated sequences were then aligned to a database of the six human proteins with BLASTP. The gene sequence and starting amino-acid position a fragment aligned to were documented and considered as a unique identifier. Paired sequences that revealed both fused fragments were in-frame with a known protein were then carried forward. This process was repeated for all replicates in the analysis and the results combined in a table of counts for each unique fragment pair in each replicate. We observed a total of 146,531 unique fragment pairs (distinct protein/amino acid start point combinations) detected in any of the replicates of the ORF selected libraries and 10,564 in replicates of the non-ORF selected libraries. The ORF selected libraries had approximately 120,000 paired-end, in-frame read counts per replicate distributed across the 146,531 unique fragment pairs. The non-ORF selected libraries had approximately 6,500 paired-end, in-frame read counts per replicate distributed across the 10,564 unique fragment pairs.

Differential growth in the higher concentrations of 3-AT versus 0 mM 3-AT is an indication of a potential protein interaction. Fragments pairs were then tested for a statistically significant increase in the number of read counts, after normalization, in the 2 and 5 mM 3-AT replicates using DESeq2 ^13^. A fold-change cutoff (based on read counts) of at least 3 and a false-discovery rate of less than 5% was applied. As expected, the positive control (Gal11p-LGF2) showed a highly significant increase in read counts in selective media (Table 1 and 2).

**Table 1.**
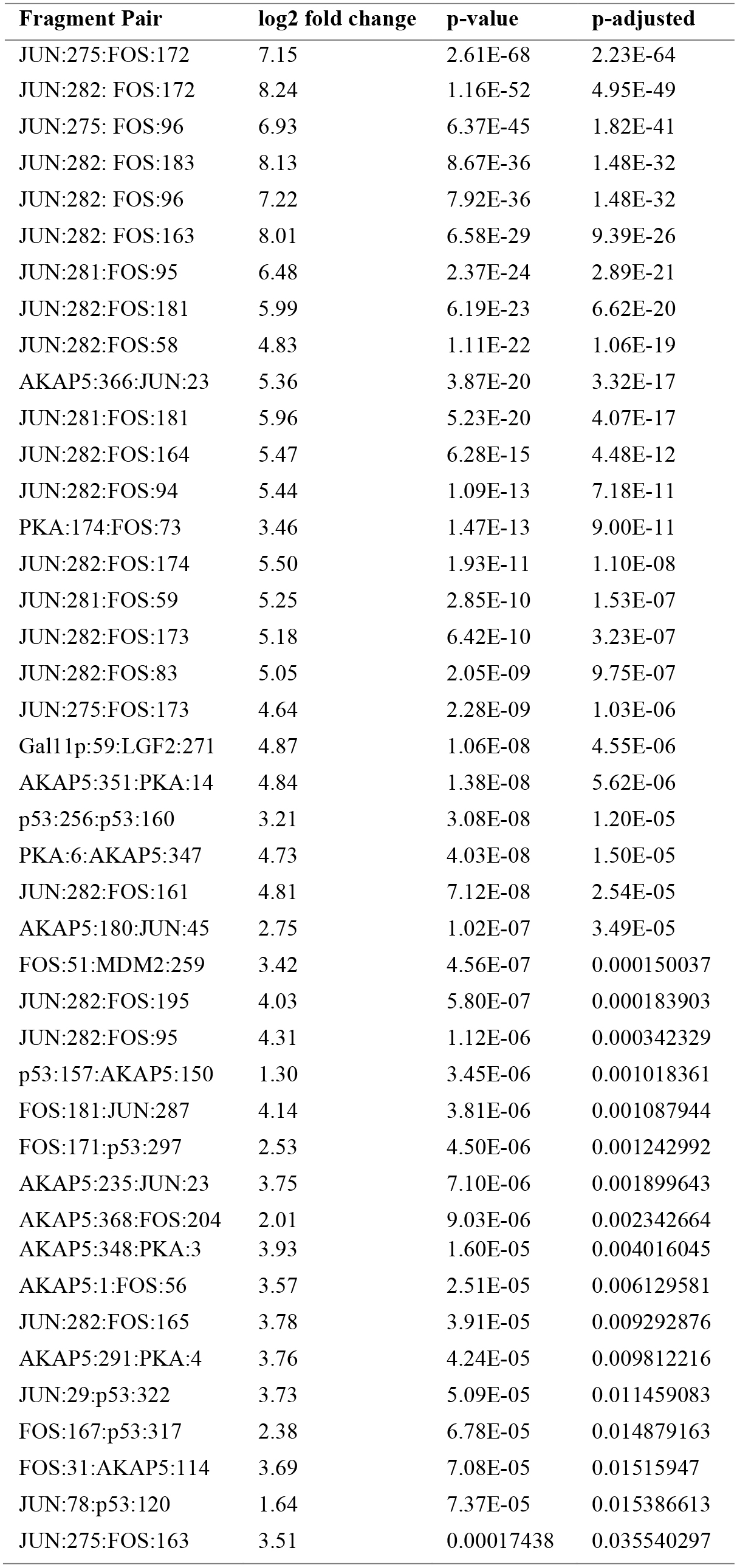
Significant interaction pairs of ORF filtered fragments 2 mM vs 0 mM 3-AT. Significantly interacting ORF filtered fragment pairs are listed with the gene name of protein 1: starting amino acid of the fragment: gene name of protein 2: starting amino acid of the fragment. The first protein in a fragment pair was fused to DBD while the second to AD. Only fragment pairs that show a positive log2 fold change with a p-adjusted (FDR) < 0.05 in presence of 2 mM 3-AT when compared to 0 mM 3-AT were deemed as significantly interacting.

| Fragment Pair | log2 fold change | p-value | p-adjusted |
| --- | --- | --- | --- |
| JUN:275:FOS:172 | 7.15 | 2.61E-68 | 2.23E-64 |
| JUN:282: FOS:172 | 8.24 | 1.16E-52 | 4.95E-49 |
| JUN:275: FOS:96 | 6.93 | 6.37E-45 | 1.82E-41 |
| JUN:282: FOS:183 | 8.13 | 8.67E-36 | 1.48E-32 |
| JUN:282: FOS:96 | 7.22 | 7.92E-36 | 1.48E-32 |
| JUN:282: FOS:163 | 8.01 | 6.58E-29 | 9.39E-26 |
| JUN:281:FOS:95 | 6.48 | 2.37E-24 | 2.89E-21 |
| JUN:282:FOS:181 | 5.99 | 6.19E-23 | 6.62E-20 |
| JUN:282:FOS:58 | 4.83 | 1.11E-22 | 1.06E-19 |
| AKAP5:366:JUN:23 | 5.36 | 3.87E-20 | 3.32E-17 |
| JUN:281:FOS:181 | 5.96 | 5.23E-20 | 4.07E-17 |
| JUN:282:FOS:164 | 5.47 | 6.28E-15 | 4.48E-12 |
| JUN:282:FOS:94 | 5.44 | 1.09E-13 | 7.18E-11 |
| PKA:174:FOS:73 | 3.46 | 1.47E-13 | 9.00E-11 |
| JUN:282:FOS:174 | 5.50 | 1.93E-11 | 1.10E-08 |
| JUN:281:FOS:59 | 5.25 | 2.85E-10 | 1.53E-07 |
| JUN:282:FOS:173 | 5.18 | 6.42E-10 | 3.23E-07 |
| JUN:282:FOS:83 | 5.05 | 2.05E-09 | 9.75E-07 |
| JUN:275:FOS:173 | 4.64 | 2.28E-09 | 1.03E-06 |
| Gal11p:59:LGF2:271 | 4.87 | 1.06E-08 | 4.55E-06 |
| AKAP5:351:PKA:14 | 4.84 | 1.38E-08 | 5.62E-06 |
| p53:256:p53:160 | 3.21 | 3.08E-08 | 1.20E-05 |
| PKA:6:AKAP5:347 | 4.73 | 4.03E-08 | 1.50E-05 |
| JUN:282:FOS:161 | 4.81 | 7.12E-08 | 2.54E-05 |
| AKAP5:180:JUN:45 | 2.75 | 1.02E-07 | 3.49E-05 |
| FOS:51:MDM2:259 | 3.42 | 4.56E-07 | 0.000150037 |
| JUN:282:FOS:195 | 4.03 | 5.80E-07 | 0.000183903 |
| JUN:282:FOS:95 | 4.31 | 1.12E-06 | 0.000342329 |
| p53:157:AKAP5:150 | 1.30 | 3.45E-06 | 0.001018361 |
| FOS:181:JUN:287 | 4.14 | 3.81E-06 | 0.001087944 |
| FOS:171:p53:297 | 2.53 | 4.50E-06 | 0.001242992 |
| AKAP5:235:JUN:23 | 3.75 | 7.10E-06 | 0.001899643 |
| AKAP5:368:FOS:204 | 2.01 | 9.03E-06 | 0.002342664 |
| AKAP5:348:PKA:3 | 3.93 | 1.60E-05 | 0.004016045 |
| AKAP5:1:FOS:56 | 3.57 | 2.51E-05 | 0.006129581 |
| JUN:282:FOS:165 | 3.78 | 3.91E-05 | 0.009292876 |
| AKAP5:291:PKA:4 | 3.76 | 4.24E-05 | 0.009812216 |
| JUN:29:p53:322 | 3.73 | 5.09E-05 | 0.011459083 |
| FOS:167:p53:317 | 2.38 | 6.78E-05 | 0.014879163 |
| FOS:31:AKAP5:114 | 3.69 | 7.08E-05 | 0.01515947 |
| JUN:78:p53:120 | 1.64 | 7.37E-05 | 0.015386613 |
| JUN:275:FOS:163 | 3.51 | 0.00017438 | 0.035540297 |

**Table 2.**
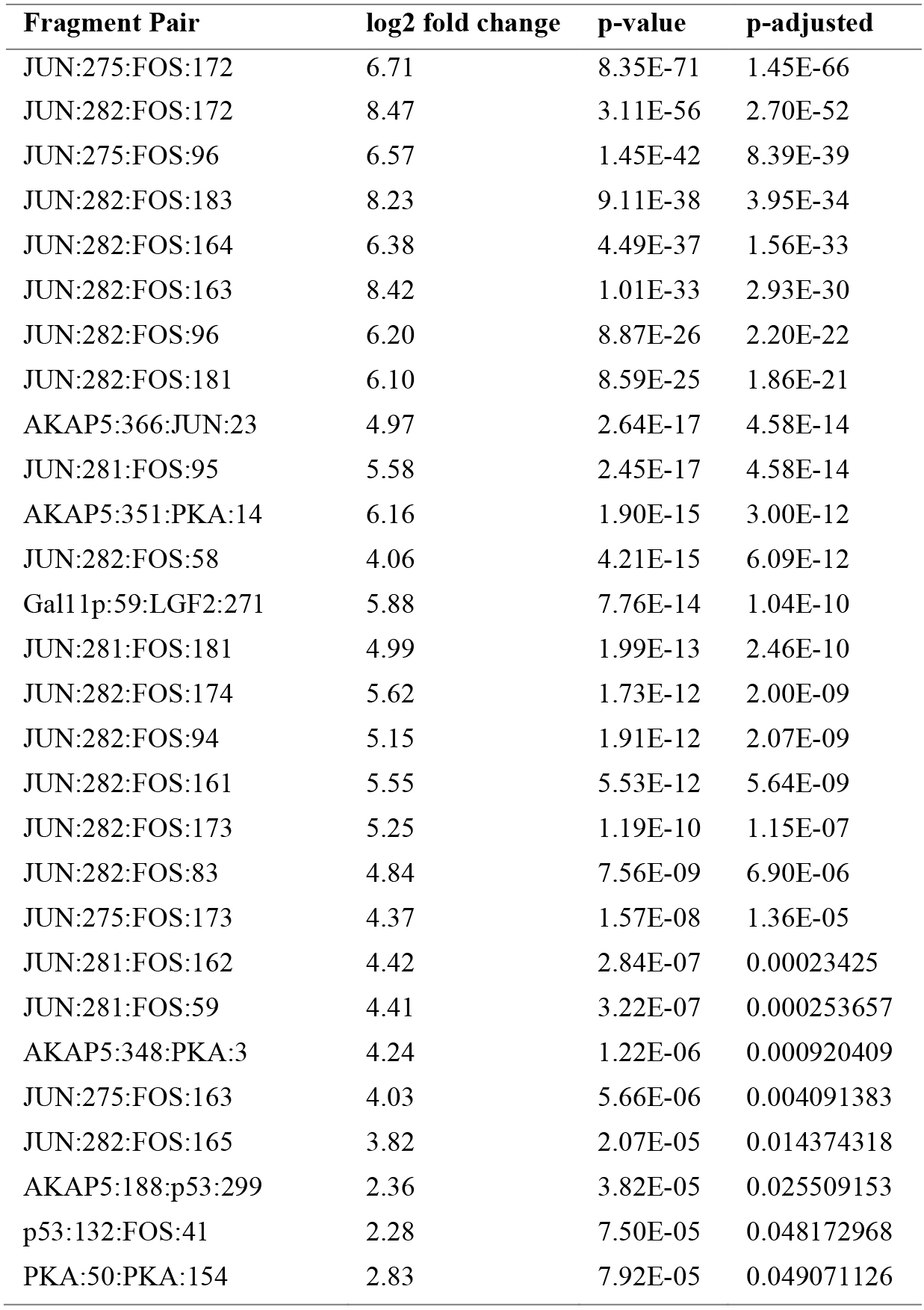
Significant interaction pairs of ORF filtered fragments 5 mM vs 0 mM 3-AT. Significantly interacting ORF filtered fragment pairs are listed with the gene name of protein 1: starting amino acid of the fragment: gene name of protein 2:starting amino acid of the fragment. The first protein in a fragment pair was fused to DBD while the second to AD. Only fragment pairs that show a positive log2 fold change with a p-adjusted (FDR) < 0.05 in presence of 5 mM 3-AT when compared to 0 mM 3-AT were deemed as significantly interacting.

| Fragment Pair | log2 fold change | p-value | p-adjusted |
| --- | --- | --- | --- |
| JUN:275:FOS:172 | 6.71 | 8.35E-71 | 1.45E-66 |
| JUN:282:FOS:172 | 8.47 | 3.11E-56 | 2.70E-52 |
| JUN:275:FOS:96 | 6.57 | 1.45E-42 | 8.39E-39 |
| JUN:282:FOS:183 | 8.23 | 9.11E-38 | 3.95E-34 |
| JUN:282:FOS:164 | 6.38 | 4.49E-37 | 1.56E-33 |
| JUN:282:FOS:163 | 8.42 | 1.01E-33 | 2.93E-30 |
| JUN:282:FOS:96 | 6.20 | 8.87E-26 | 2.20E-22 |
| JUN:282:FOS:181 | 6.10 | 8.59E-25 | 1.86E-21 |
| AKAP5:366:JUN:23 | 4.97 | 2.64E-17 | 4.58E-14 |
| JUN:281:FOS:95 | 5.58 | 2.45E-17 | 4.58E-14 |
| AKAP5:351:PKA:14 | 6.16 | 1.90E-15 | 3.00E-12 |
| JUN:282:FOS:58 | 4.06 | 4.21E-15 | 6.09E-12 |
| Gal11p:59:LGF2:271 | 5.88 | 7.76E-14 | 1.04E-10 |
| JUN:281:FOS:181 | 4.99 | 1.99E-13 | 2.46E-10 |
| JUN:282:FOS:174 | 5.62 | 1.73E-12 | 2.00E-09 |
| JUN:282:FOS:94 | 5.15 | 1.91E-12 | 2.07E-09 |
| JUN:282:FOS:161 | 5.55 | 5.53E-12 | 5.64E-09 |
| JUN:282:FOS:173 | 5.25 | 1.19E-10 | 1.15E-07 |
| JUN:282:FOS:83 | 4.84 | 7.56E-09 | 6.90E-06 |
| JUN:275:FOS:173 | 4.37 | 1.57E-08 | 1.36E-05 |
| JUN:281:FOS:162 | 4.42 | 2.84E-07 | 0.00023425 |
| JUN:281:FOS:59 | 4.41 | 3.22E-07 | 0.000253657 |
| AKAP5:348:PKA:3 | 4.24 | 1.22E-06 | 0.000920409 |
| JUN:275:FOS:163 | 4.03 | 5.66E-06 | 0.004091383 |
| JUN:282:FOS:165 | 3.82 | 2.07E-05 | 0.014374318 |
| AKAP5:188:p53:299 | 2.36 | 3.82E-05 | 0.025509153 |
| p53:132:FOS:41 | 2.28 | 7.50E-05 | 0.048172968 |
| PKA:50:PKA:154 | 2.83 | 7.92E-05 | 0.049071126 |

Analysis of read counts for fragment pairs with at least one observed read in the libraries with selection versus 0 mM 3-AT libraries showed a correlation (r^2^) of 0.67, 0.97 and 0.97 for 0, 2, and 5 mM replicates, respectively (Sup. Fig. 2). The lower correlation among the 0 mM replicates is likely due to the fragment pairs with significant numbers of reads in 2 mM or 5 mM 3-AT had very few reads in 0 mM 3-AT making correlation less likely. We observed a correlation of -0.09 and -0.07 when 0 mM 3-AT libraries were compared to 2 mM and 5 mM 3-AT, respectively. Comparison of a library with no carrier (no DMSO, no 3-AT) vs the 0 mM 3-AT library showed correlation of 0.64, agreeing well with the correlation among 0 mM 3-AT replicates. This suggests that the 0 mM 3-AT selected libraries do not appear to contain bias from the DMSO carrier. Whether the growth in minimal media creates an initial selective pressure that could result in bias of fragment content is not clear but all comparisons in this study were controlled with 0 mM growth levels.

### Protein Interaction Analysis

Using this method, we have tested at high-resolution 96.14% (5686520/5914624 pairwise amino acid combinations) of the possible interaction space when all libraries are considered (Fig. 3a). This was possible by generating protein fragments with multiple starting points. The tested interaction space was reduced to 5.6% (331580/5914624) and 1.9% (111597/5914624) of significantly interacting amino acids in the 2 mM and 5 mM concentrations of the 3-AT, respectively. Figure 3b and c show the refined interaction space of the 36 protein pairs (6 proteins x 6 proteins) used in this study. By looking at the change in interaction regions moving from 2 mM 3-AT (Fig. 3b) to an increase in selective pressure using 5 mM 3-AT (Fig. 3c) there are obvious changes in the interaction landscape. These data indicate the depth of information generated using mild and strong selective pressure (correlating, in theory, to mild and strong interactions).

**Figure 3.**
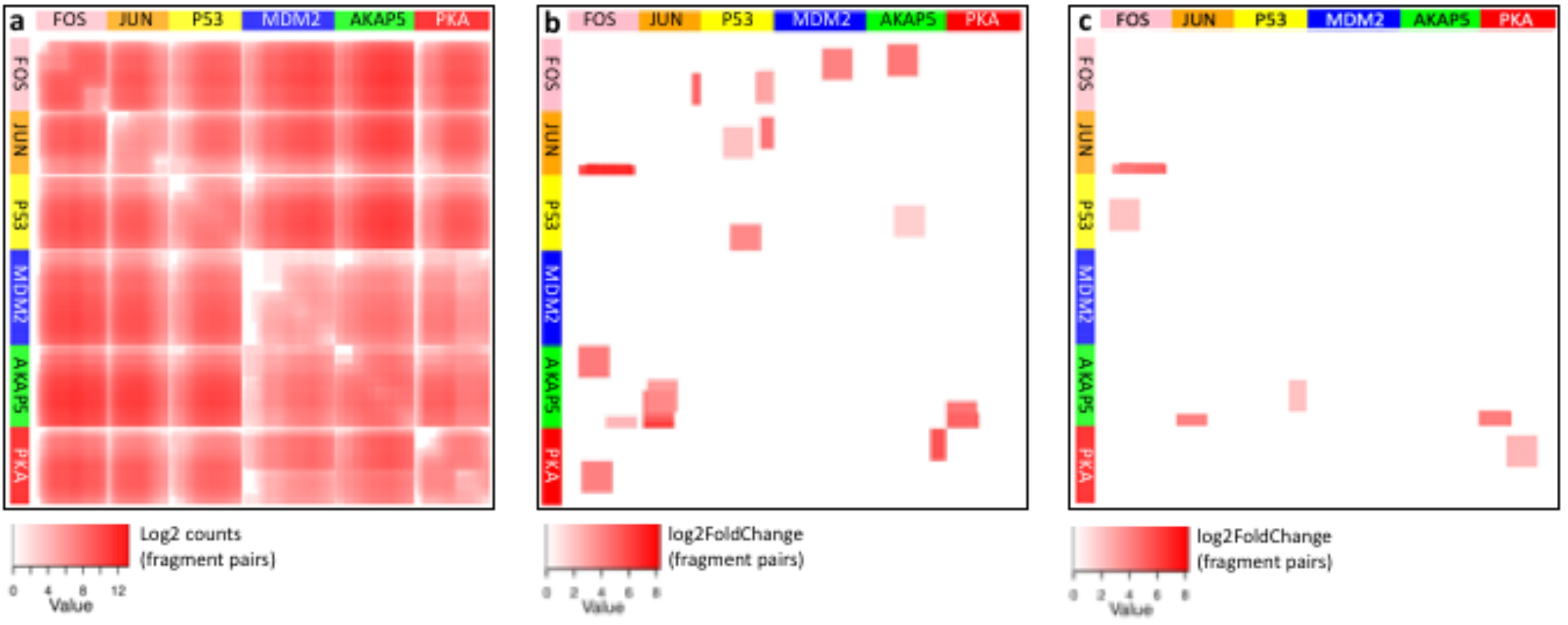
Matrix heatmaps of all Amino Acid (AA) pairs tested or interacting in this study. Colored panels on the top and left show location of proteins in the matrix. (**a**) Detection of the AA pair (red) or no detection of the AA pair (white) in any of the libraries sequenced. (**b)** and (**c)** show log2 fold change in 2 mM and 5 mM libraries with respect to the 0 mM library for fragment pairs that show a statistically significant enrichment. Enriched pairs in (**b**) and (**c**) imply interaction between the corresponding regions of the proteins. Orientation of the tested pair is included to illustrate not all interactions are symmetrical in the matrix, i.e. fusions to DBD are listed vertically while fusions to AD are horizontal.

Figure 4 illustrates several of the most significant (p-adjusted < 0.05) fragment pairs of the expected interactions (Tables 1 and 2). The JUN-FOS protein fragments demonstrated the most abundant interaction out of the 36 possible interaction combinations (not including the different starting amino acids) comprising 23/42 and 21/28 significant interactions in 2 and 5 mM, respectively (Tables 1 and 2). Additionally, in the non-ORF filtered data this interaction was 35/38 and 28/31 significant interactions in 2 and 5 mM, respectively (Sup. Tables 2 and 3). Importantly, the fragments align to the interaction regions for the proteins (Fig. 4, light grey) and show fragments in both orientations demonstrating the level of detail this method affords (Tables 1 and 2). The JUN-FOS interaction also showed similar fragment pairs for the ORF filtered and non-ORF filtered libraries which also attests to the strength of the interaction. The multiple interactions of JUN and FOS fragments are consistent with the interaction sites reported in literature ^6,14^.

**Figure 4.**
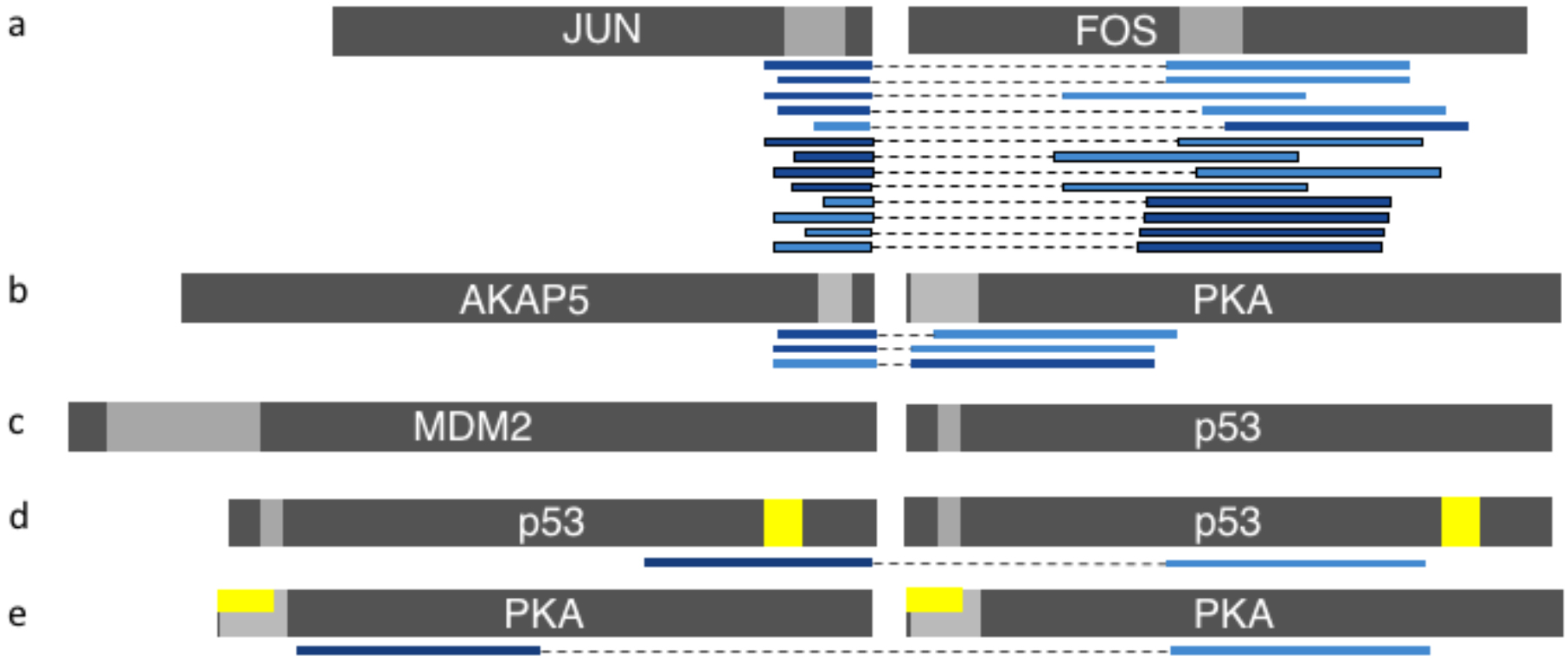
Significant interactions detected (p < 0.05) for expected protein pairs. (**a**) interacting pairs between JUN-FOS. (**b**) interacting pairs between AKAP5-PKA. (**c**) expected interaction regions of MDM2 and p53. (**d**) p53 homodimer and (**e**) PKA homodimer. The light grey region of the protein schematic indicates interaction regions reported in the literature ^6,9,11,14,15^. ORF filtered fragments are shown with no outline and the non-ORF filtered fragments have a black outline. Dark blue fragments are associated with the activation domain (AD) and the light blue fragments are associated with the DNA binding domain (DBD). Yellow bars indicate residues involved in homodimer association.

In addition, Fig. 4b shows AKAP5 and PKA also had interacting fragment pairs in both orientations. For the ORF filtered fragments, the significant interactions for AKAP5 and PKA comprised 4/23 and 2/28 significant interactions in 2 and 5 mM, respectively (Tables 1 and 2). An advantage to this fragment-based method, rather than more traditional full-length protein, is the ability to identify the interaction region(s) between the proteins. Looking at Fig. 4 b more closely shows alignment of the significant interaction pairs corresponds to the interaction regions described in the literature for AKAP5 and PKA as well ^9^.

Although the literature shows p53 and MDM2 interact, the AVA-seq method was not able to detect statistically significant interactions between the two proteins (Fig. 4c). This is likely due to the complex nature of the interaction. The interaction site of MDM2 is large, and the interaction complex of MDM2-p53 is dependent primarily on van der Waals which is different from most identified proteins. Furthermore, interaction occurs in a buried surface that consists mostly of hydrophobic interaction between pseudosymmetry domains ^11,15^. Fragments from these proteins were however detected as significant interactions with other proteins (Sup. Fig. 3). Interestingly, there was a significant self-interaction between p53-p53 and PKA-PKA (Fig. 4d and e, respectively). The yellow bars in Fig. 4d-e represent residues involved in the p53-p53 dimer interface ^16^ or PKA-PKA dimer interface ^17^. Two of the four fragments involved in the self-interaction align to these regions involved in the dimer interaction.

This method not only identified known self-dimerization interactions (Fig. 4 d-e) it also shines light on possible novel interactions which have not been explored in the literature to the best of our knowledge (Sup. Fig. 3, Tables 1 and 2, Sup. Tables 2 and 3). Several of the interactions indicate fragments in both orientations, increasing the probability of a real, strong interaction and worth investigating further. The AVA-seq method is able to confirm at high resolution the interactions of 2 of 3 of the human proteins pairs as well as two homo-dimer interactions and other, possibly novel, interactions between the 6 proteins.

## Discussion

Protein-protein interaction data can provide important information for understanding how an individual protein functions and its system-wide role in the context of other proteins. Despite the importance, methods such as the two-hybrid system have not seen significant reductions in labor and cost since its first use. Though some improvements have been made using next-generation sequencing, deep screening or large-scale two-hybrid studies remain labor intensive.

At the center of our approach to utilizing NGS for two-hybrid based protein-interaction mapping is the convergent fusion vector, pAVA. The novelty of pAVA is it joins the traditionally individual “bait” and “prey” DNA sequences on a single DNA molecule allowing them to be amplified and paired-end sequenced. This combined with the high transformation efficiency has allowed us to test almost the entire interaction space of the 6 proteins at high resolution (146,000 paired fragments tested). As with methods such as RNA-seq, this system is limited simply by diversity of the library and depth of sequencing. Higher resolution of interacting domains is achievable with deeper sequencing of diverse libraries. An important feature of the system is the ability to “dial-in” the level of selection by changing the concentration of 3-AT. Here, we used 2 and 5 mM concentrations but this could be changed depending on the strength of the interactions being targeted. Our results show that many of the interactions detected in the more stringent 5 mM selection are also found in the 2 mM 3-AT selection, while some interactions are only detected in one of the conditions. It may be possible, in the future, to rank the strength of interactions in the system relative to the positive control (Gal11p-LGF2) as it is included in every experiment. Here, for example, the JUN-FOS fragment interactions in general were stronger relative to the positive control in the 2 mM 3-AT selection while the PKA-AKAP5 fragments were weaker.

We observed 2 of 3 known interactions among the 6 proteins tested. Those are the JUN-FOS and PKA-AKAP5 interactions. Homodimerization of p53 and PKA were also observed, albeit by one fragment pair each (Fig. 4d-e). The p53-MDM2 interaction was not detected at a statistically significant level by the system showing its limitation in detecting interactions between large domains (Fig. 4c). Frequent criticism of two-hybrid systems centers around the potential for false-positives, or interactions that should not be detected; and false-negatives, that is, interactions that were missed. The AVA-seq system employs various features to mitigate both types of errors. While this does not guarantee that the interactions detected occur *in vivo*, it increases the likelihood that interactions are not spurious within the system or were simply missed. These features include interactions being observed in both orientations, that is with the bait and prey fragments fused to the DBD and AD and vice-versa. Additionally, applying a requirement that multiple, overlapping fragments from the same genes be observed to interact decreases the possibility that the interaction is invalid. Removal of auto-activators, those fragments that activate the system regardless of which protein fragment it is paired with, can be removed by applying an upper bound cutoff to the number of fragments it is reported to interact with. While current limitations of NGS fragment lengths require that in general we test fragments rather than full-length genes, we observed benefits from testing multiple fusions at various amino acids positions for each protein. We noted that not all fragment pairs expected to interact were observed and that likely not every fusion point creates a functionally active protein to allow interaction ^12^. Thus, using multiple fusion points within a gene, rather than a full-length gene, may help overcome both false-positives and false-negatives. However, the limitation of fragment length was clear in the missed p53- MDM2 interaction likely due to larger fragments being required to span the interaction region ^11^. Lastly, the ability to screen with multiple levels of selective pressure in a cost-effective manner should increase the chances of detecting a range of interactions.

The reduction of the potential interaction space by approximately 17-fold (2 mM) and 50-fold (5 mM) to a small portion of interactors is significant. It is even more significant a reduction when considering that 3 pairs of the proteins were expected to interact. The additional interactions detected in this study (Sup. Fig. 3) provide a starting point for future validation studies. Some of these interactions such as that between AKAP5 and p53 show interaction between multiple overlapping fragment pairs with at least one in the reverse-fusion orientation.

With the improvements we have made to the two-hybrid system we envision two uses for AVA-seq. The first, as demonstrated here, is the high resolution protein-protein interaction mapping of a small set of proteins either all-versus-all, or few-versus-all. This might include proteins of unknown function being tested against a whole cDNA library or those from a single pathway. The number of transformants and sequencing depth is readily achievable for high resolution domain mapping. The second application will expand to large all-versus-all interaction screening for entire bacterial genomes and beyond. By utilizing the ORF selection process, the number of screening events have been drastically reduced while maintaining the benefits of fragment-based mapping described above. Indeed, using the high transformation efficiency of a single vector and deep sequencing we believe whole-genome protein-interaction mapping could be achieved by small laboratory groups in a relatively short amount of time.

## Online Methods

### Design of pBORF vectors

The pBORF plasmid was designed by replacing the AmpR in the pBlueScript II SK (+) vector (Stratagene #212205) with a kanamycin resistance gene just after the AmpR promoter. The β-lactamase localization sequence and the enzyme-coding portion of β-lactamase were inserted in the multiple cloning site region of the modified pBlueScript II SK (+) vector. Approximately 25 bp inserts were designed and inserted between the β-lactamase localization sequence and the β-lactamase gene to allow for ORF filtering and to differentiate pBORF associated with DBD (λcI), referred to as pBORF-DBD, from the pBORF associated with AD (RNAP), referred to as pBORF-AD. Using the modified pBlueScript vector from above as a template, primer A and primer B were used to create an insert for pBORF-AD and primer C and primer D to create an insert for pBORF-DBD. Primers were subjected to annealing program (95 °C for 2 min, slow cool to 25 °C) then diluted 1:1000 from 50 μM to 50 nM. To prepare for ligation, the modified pBlueScript plasmid was linearized with primers E and F. Ligation of the linearized plasmid was setup using 3:1 insert to vector where the insert. The resulting ligations created pBORF-DBD or pBORF-AD.

When needed, pBORF-DBD and pBORF-AD were linearly amplified the same day as ligation. The PCR used primers G and H for the pBORF-AD plasmid and primers G and I for the pBORF-DBD plasmid. The column purified linearized pBORF-DBD and -AD vectors were then subjected to Dpn1 (NEB; R0176S) digestion followed by another PCR column cleanup.

### Design of pAVA

The pAVA plasmid was constructed from the BacterioMatch II two-hybrid system (AGILENT) vectors pBT and pTRG. First, the AD domain in pTRG was amplified using primers that included XhoI and NotI restriction sites. Restriction digestion was performed for both pBT plasmid and the amplified AD PCR product using XhoI and NotI enzyme sites. Next, ligation was performed and resulted in the AD in convergent orientation with DBD in pBT plasmid. This new construct will be referred to as pAVA (plasmid all-versus-all). The pAVA plasmid was linearized using primers N and O. Column cleanup was performed after amplification using GenElute PCR Cleanup Kit then subjected to DpnI treatment using standard protocol. To introduce BstX1 restriction enzyme sites, primers P and Q were used. Ligation was performed with 3:1 insert to vector ratio using linearized pAVA plasmid digested with BstX1 and BstX1 digested insert obtained from P and Q primer amplification.

### pAVA controls

Gal11p and LGF2 sequences (sp|P04386|GAL4_YEAST and sp|P19659|MED15_YEAST, respectively) were amplified from the pBT-LGF2 and pTRG-Gal11p control plasmids from the BacterioMatch II Two-Hybrid System and ligated into both pBORF-AD and pBORF-DBD. The final converging positive control constructs in pAVA were made as described above (Fig. 2b-c). The first negative control is the empty plasmid containing AD and DBD without DNA insert (Fig. 2a). The second and third negative controls are the positive control constructs with the addition of one nucleotide to introduce a frame shift to LGF2 only or both LGF2 and Gal11p (Fig. 2d-e). Each control plasmid was tested separately in the presence of 0, 2, and 5 mM 3-amino-1, 2, 4-triazole (3-AT) in DMSO as well as unselected sample which includes no DMSO or 3-AT as an additional control.

### Amplification of 6 Human Genes

The following 6 human genes were purchased from Origene as 10 μg stocks in pCMV6 Entry vector with Sgf1 and Mlu1 cloning sites: PKAR2 (RC220376, 1212 bp), AKAP5 (RC221314, 1281 bp), MDM2 (RC219518, 1491 bp), p53 (RC200003, 1179 bp), FOS (RC202597, 1140 bp) and JUN (RC209804, 993 bp). All sequences were confirmed by in-house Sanger sequencing. Each gene in pCMV6 Entry was amplified separately using T7 and M13 reverse primers.

### Shearing of DNA and Ligation of Fragmented Genes into pBORF Vectors

30 nM of PKA, AKAP5, MDM2, p53, FOS and JUN were pooled in a total volume of 50 μL and transferred to a Covaris microTUBE (Covaris, Part No. 520045, Lot No. 002563). The sample was sheared using the settings appropriate for 400 bp median size. Next, end repair (NEBNext DNA Library Prep Master Mix Set for Illumina, NEB, Cat. No. E6040S) was performed as directed. Blunt ligation of the end-repaired fragments into linearized pBORF-DBD and pBORF-AD vectors were performed using molar ratio of 6:1 (insert to vector).

### Open Reading Frame (ORF) Filtering

Transformations were performed for each pBORF-AD and pBORF-DBD ligation products using CopyCutter EPI400TM Electrocompetent *E. Coli* (Lucigen Cat. No. C400EL10; 1.8 mV). The transformations for pBORF-DBD or pBORF-AD were pooled. In order to have an idea of how many unique fragments are being represented in the library, dilutions were made and plated on LB-agar supplemented with KAN (30 μg/mL) allowing for the total number of colonies to be counted. The colonies on the KAN antibiotic alone represent the non-ORF fragments. Transformed cells plated directly on LB-agar plates with KAN (30 μg/mL) and CB (15 μg/mL) represent the ORF filtered fragments. We typically see a 95-99% decrease in survival when comparing the number of non-ORF : ORF fragments.

### Extracting DNA

Colonies grown on KAN were gently scraped and resuspended with sterile LB. The same was done for the colonies grown on KAN and CB. ~100 μL of the resuspended colonies were pelleted and DNA was extracted with a GenElute Plasmid MiniPrep (Sigma, Cat. No. PLN350). Samples were then subjected to 0.7% agarose gel (110 V, 1 h) and gel purified.

### Extracting ORF Filtered DNA Product from the pBORF Vectors: Overlap Extension PCR

Extracting the ORFs from the vector was carried out by PCR. PCR amplification was performed using ORF filtered DNA in pBORF-DBD or pBORF-AD vectors and primer J and K for pBORF-DBD or primer L and M for pBORF-AD. Next, overlap extension PCR was performed using the resulting products above plus primers J and L. The PCR cycle was as follows: 98 °C for 2 min, 12x(98 °C 15 s, 57 °C 20 s, 72 °C 45 s), 72 °C 3 min, 4 °C hold. The resulting PCR product combined ORF filtered pBORF-DBD and pBORF-AD fragments into one larger product. This whole process was repeated for the non-ORF using the same protocol.

### Ligation of PCR product into pAVA

The gel purified pAVA was linearized with BstX1 using a standard protocol. Next the samples were run on a 0.7% agarose gel, the band was excised and extracted GenElute Gel Extraction kit. The PCR product from the overlap extension PCR was digested with BstX1 in the same manner as above. Next the samples were run on a 1.5% agarose gel and the bands between 700-1200 bp were excised and extracted.

The gel purified BstX1 digested products were ligated with a 1:6 vector to insert ratio with T4 DNA Ligase (NEB) overnight followed by GenElute PCR cleanup. The purified ligation were then transformed into NEB Turbo Electrocompetent cells and plated on LB-agar plates supplemented with 10 μg/mL CAM (chloramphenicol). The colonies are then scraped with LB as described above and DNA was then extracted. DNA was transformed into the BacterioMatch II Electrocompetent Reporter Cells (Stratagene, Cat. No. 200195).

### 3-AT selection

The resulting colonies in the BacterioMatch II Reporter strain are scraped with LB broth. 500 μL of the slurry are diluted into 5 mL of minimal media and centrifuged (5 min, 3,000 x g, RT). The supernatant was decanted and the pellet was washed with another 5 mL minimal media. This wash of the pellet was repeated for a total of 4 washes followed by resuspension in 5 mL fresh minimal media. The OD_600_ of the cells was measured and diluted to OD_600_ = 0.05 in 75 mL minimal media.

A 5 mL culture of the Gal11p-LGF2 (positive control in pAVA and VR) was inoculated with fresh colonies and grown in LB broth in the presence of 25 μg/mL CAM for 3 hours. The sample was then centrifuged (5 min, 3,000 x g RT) and washed identically to the slurry above. The final resuspension for the positive control was OD_600_ = 0.05 in 1 mL minimal media. From this a 1:1000 dilution was made in 1 mL of minimal media. 7.5 μL was removed from the 75 mL cell mixture of the washed sample and replaced with 7.5 μL of the 1:1000 dilution of the positive control for a final OD_600_ = 5×10^-9^ spike in.

One 15 mL culture tube was set up for an unselected sample that contained 5 mL of cells (OD_600_ = 0.05 with the positive control spike in) in minimal media. Three additional tubes with 5 mL of cells for each of the following at 0 mM (25 μL DMSO), 2 mM (15 μL DMSO and 10 μL 1 M 3-AT) and 5 mM 3-AT (25 μL 1 M 3-AT) for a total of 10 culture tubes. Cells were allowed to grow for 9 h, 250 RPM, 37 °C. After 9 h of growth, OD_600_ was measured. Samples were centrifuged at 3,000 g for 5 min. DNA was extracted using GenElute MiniPrep.

### Library Construction

To begin library construction, interacting fragments from the 3-AT selection were amplified from the pAVA plasmid using primers R and S (Sup. Table 1). The sequencing libraries are prepared using NEBNext Ultra II DNA Library prep kit (Cat. No. E7645S) according to the manufacturer’s protocol. The library concentrations are determined using KAPA Library Universal Quantification Kit (KK4827). Agilent High Sensitivity DNA Kit (5067-4626) for Bioanalyzer electrophoresis was used to determine average fragment length. Final libraries were prepared as directed in the MiSeq System Denature and Dilute Libraries Guide (support.illumina.com) and run using MiSeq V2 Reagent Kit (300 Cycles) (MS-102-2002).

### Preparation of media and reagents for selection 3-AT selection

For each selection experiment, 100 mL of fresh minimal medium was made using the following recipe adapted from the BacterioMatch II system manual. In the following order, 2 mL of 20% glucose, 1 mL 20 mM adenine HCl and 10 mL 10x His-dropout amino acid mix (Clontech, 630415, autoclaved according to manufacturer’s directions, stored at 4 °C until use) were combined. Then 100 μL of each of the following: 1 M MgSO_4_, 1 M Thiamine-HCl, 10 mM ZnSO_4_, 100 mM CaCl_2_ and 50 mM IPTG were added. After mixing well, 76 mL autoclaved millipure water, 10 mL 10x M9 salts and 100 μL of 25 mg/mL CAM were added. To make 10x M9 salts, dissolve 14 g disodium phosphate (Na_2_HPO_4_*7H_2_O), 6 g potassium dihydrogen phosphate (KH_2_PO_4_), 1 g sodium chloride (NaCl) and 2 g ammonium chloride (NH_4_Cl) in 200 mL millipure water, filter and autoclave. Store at room temperature. 1 M 3- AT stocks were made in 100% DMSO and stored at −20°C for no more than 30 days. Adenine, IPTG, Thiamine-HCl and 3-AT aliquots were used once, and any remaining was discarded.

### Analysis of significant interactions

A higher growth in the presence of 3-AT at 2 mM and 5 mM concentrations versus 0 mM is indication of a potential protein-protein interaction. Statistical significance of differential growth for each comparison was evaluated from three replicates in each growth condition at a positive predictive value of 95% (FDR < 0.05) using DESeq2 ^13^. For each comparison, only those protein fragment pairs that were observed in at least 4 of the samples out of 6 replicates were taken for differential growth analysis to estimate the significant differences. DESeq2 performs an internal normalization step where the geometric mean is calculated for each row across all samples and counts in each sample are then divided by the mean. The median of the ratios in a sample is used as the size factor for that sample to correct for the library size and composition bias. Rows containing count outliers are automatically removed using Cooks’s distance. In addition, an optimization procedure further removes the fragment pairs with low counts by filtering the rows where mean of normalized counts is below a determined threshold. Finally, a negative binomial generalized linear model is fitted to determine differential growth using the Wald test for significance testing which computes p-value and the adjusted p-values (FDR) for each protein fragment pair. Only fragment pairs that show a positive log2 fold change with an FDR < 0.05 in presence of 3-AT when compared to 0 mM 3-AT were deemed as significantly interacting.

## Acknowledgments

The authors would like to thank the members of the Genomics Laboratory at Weill Cornell Medicine in Qatar for their assistance. This research was supported by the BMRP grant funded by Qatar Foundation and the WCM-Q Pilot Grant.

## Author contributions

Design of the work (S.A. and J.M.), acquisition and/or analysis (all authors), interpretation of data (S.A., S.S-R., N.A., I.A. and J.M.) and drafting of the manuscript (S.A., S.S-R., N.A., I.A. and J.M.). All authors have approved the final version of this manuscript.

